# Putting causality into context: Causal capture escapes the visual adaptation of causality

**DOI:** 10.1101/2025.04.10.648104

**Authors:** Ben Sommer, Martin Rolfs, Sven Ohl

## Abstract

Results from psychophysical studies using visual adaptation suggest that launch detectors in the visual system underlie the perception of causality in simple visual events. These detectors respond to events in which one stimulus collides with another stimulus (i.e., a launch), and do not respond to events where one stimulus passes over another (i.e., a pass). Prolonged visual adaptation to launches significantly reduces observers’ propensity to see causal launches at the same retinotopic location. This finding could be taken to indicate that launch detectors are necessary for the local detection of causal launches. However, contextual events—that are spatially separated from the test event location—shift observers’ perception of a causal relation in the direction of the type of contextual event (Scholl & Nakayama, 2002), providing evidence for spatial integration beyond a specific retinotopic location. Here, we used visual adaptation as a tool to investigate whether the contextual influence on causal perception relies on local launch detectors. Before and after adaptation, we determined the proportion of reported launches in ambiguous test events in the presence of no context, launch context, and pass context events. We hypothesized that if the contextual influence relies on (unadapted) local launch detectors, then visual adaptation should affect the contextual influence on causal perception. Before adaptation, a launch-context event increased the proportion of reported launches (while a pass context event decreased it). Visual adaptation to launches significantly decreased the proportion of reported launches in no-context trials, but did not affect perceptual reports in no-context trials. In fact, contextual influences, expressed relative to no-context trials, emerged strongly after adaptation. This result suggests that context effects override strong negative aftereffects from adaptation, indicating that contextual influences operate at a level that bypasses the local launch detector at the adapted location.

## Introduction

Cause and effect underlie physical interactions. For an agent, understanding the systematic causal relations provides an advantage for successfully acting in its environment. For instance, understanding that pushing against an object changes the object’s position allows the agent to open a door or to get an obstacle out of its way. Humans report a vivid causal impression when perceiving events in which one object’s motion cooccurs in space and time with the onset of another object’s motion (e.g., one moving object is seen to launch the movement of a second object when it starts to move right after contact with the first; Michotte, 1963). The causal impression in such launching events occurs fast, and automatic, and it is irresistible (Michotte, 1963; Scholl & Tremoulet, 2000). Moreover, the causal impression depends on the exact spatiotemporal parameters of the event. Either delaying the movement of the effect object after the contact or introducing spatial overlaps or gaps reduces the causal impression. These findings suggest that the visual system is host to finely tuned launch detectors that underlie the perception of causal relations (Michotte, 1963; Scholl & Tremoulet, 2000). Yet it remains notoriously hard to distinguish between predictions from a visual and a cognitive perspective (Rips, 2011).

Visual adaptation studies have fueled the idea that the detection of a causal relation occurs already in the visual system (Kominsky & Scholl, 2020; Ohl & Rolfs, 2025; Rolfs et al., 2013). In these studies, observers reported whether they saw a launch or a pass in ambiguous test events. Importantly, the presentation of a launch-adaptor reduced the proportion of perceived launches in subsequent trials. An intriguing aspect of that finding is that the visual adaptation occurred in a retinotopic reference frame (Kominsky & Scholl, 2020; Rolfs et al., 2013), demonstrating the involvement of retinotopically organized visual areas in perceiving causal relations. Moreover, the adaptation does not transfer across different motion directions (Ohl & Rolfs, 2025), suggesting that the perception of causality is computed in (or based on the output of) direction-selective channels in the visual system.

Whether and how we perceive causality in simple launching events is not only determined by the presence of a collision at the test event’s location but also depends on context events (Choi & Scholl, 2004, 2006; Scholl & Nakayama, 2002). When one object moves towards a stationary second object until they fully overlap, and only then the second object starts to move, observers typically report seeing a non-causal pass (i.e., the first object passes the second stationary object). However, the display of a non-causal pass can be perceived as a causal launch when a launch-context event is presented along with the non-causal pass (i.e., causal capture; Scholl & Nakayama, 2002).

In this study, we are pitting two opposing influences on causal perception against each other (i.e., visual adaptation vs. contextual events). Specifically, we asked whether interfering with the perception of causality by means of visual adaptation would interfere with the contextual influence on causal perception. To this end, we determined whether the contextual influence would be diminished after the visual adaptation at the test event location. If the adaptation directly reduces the contextual influence, then launch detectors at the adapted visual location would seem necessary for the contextual influence. In contrast, if the adaptation does not attenuate the contextual influence, then the contextual influence must be established through a separate mechanism that *bypasses* the adapted location. In line with previous findings, we observed that adaptation strongly reduced perceived causality in ambiguous test events (Kominsky & Scholl, 2020; Ohl & Rolfs, 2025; Rolfs et al., 2013). Surprisingly, we observed that adaptation at the test event location significantly enhanced causal capture.

## Method

### Participants

We recruited 11 participants who had normal or corrected-to-normal vision. In the first session, we determined if participants were able to distinguish between launches and passes by determining whether the proportion of reported launches decreased with increasing disk overlap. Two observers that did not fulfill this criterion were excluded. The final sample consisted of nine human observers (aged 19–34; 9 female, 9 right-handed, 7 right-eye dominant) that were tested in 3 sessions (1 training session without adaptation; 2 test sessions with adaptation). Data obtained in the training session did not enter the final analyses. Participants were compensated either by course credit or at a rate of 10€/hr. Before any data was collected, all participants signed an informed consent sheet and were again verbally informed of the data collection procedures, DSGVO guidelines and given a brief introduction to the experimental paradigm. The study complies with the Declaration of Helsinki (2008) and was approved by the Ethics Committee of the Department of Psychology at Humboldt-Universität zu Berlin.

### Material and Procedure

Participants sat in a sound-shielded, dimly lit room with their head on a chin rest. Their eye position was controlled for by tracking their dominant eye using an Eyelink 1000 Desktop Mount eye tracker (SR Research, Ottawa, ON, Canada) with a sampling rate of 1000 Hz. We presented the visual stimuli on a video projection screen (Celexon HomeCinema, Tharston, Norwich, UK) using a PROPixx DLP projector (VPixx Technologies Inc., Saint Bruno, QC, Canada) at a spatial resolution of 1920 × 1080 pixels and a refresh rate of 120 Hz. The screen was mounted on the wall 180 cm away from the observers. The experiment was run on a DELL Precision T7810 (Debian GNU Linux 8) and implemented in Matlab R2023b (Mathworks, Natick, MA, USA) using the Psychophysics toolbox 3 (Brainard, 1997; Kleiner et al., 2007; Pelli, 1997) for stimulus presentation and the Eyelink toolbox (Cornelissen et al., 2002) to control the eye tracker. Participants gave their behavioral responses by pressing one of two keys on a standard keyboard. In this experiment, the observers’ task was to report a test stimulus as either a launch or a pass (**Figure 1a**).

**Figure 1.**
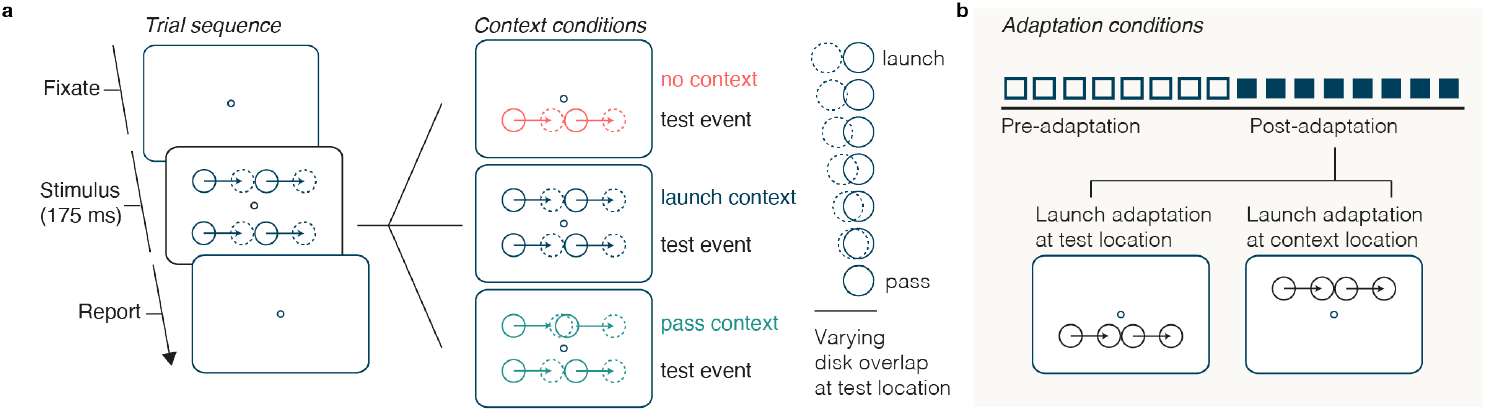
Contextual influence on the perception of causality before and after adaptation. **a** During a trial, participants fixated the center of the screen and were presented with stimuli that differed in the amount of disk overlap, ranging from 0 to 100 % in 7 equidistant steps, where 0% overlap constitutes a launch event and 100% overlap constitutes a pass event. Reports were given by pressing either the up or down arrow on a standard keyboard to indicate launch or pass. There were 3 context conditions (i.e., no context, launch context, pass context). **b** These conditions would then be tested over 16 blocks, with the second half of the blocks including an adaptation procedure for each block. For the adaptation procedure, participants were shown 320 launch events in quick succession at the start of a block and then 16 events at the start of a trial to top-up adaptation. The adaptation location could be at the test location (below fixation) or at the context location and was varied across sessions. In total, each block included 42 trials, as there were 7 disk overlaps × 3 context events × 2 repetitions.

Before testing started, participants were verbally instructed about the nature of the task. Demos of launch and pass events were shown as part of that introduction. In a trial, we presented test events of varying degrees of disk overlap. In launch events (0% overlap), the two disks touch tangentially when the second disk starts to move, and the event is typically perceived as a causal launch. In pass events (100% overlap), the disks are completely superimposed for one frame when the second disk starts to move, which participants typically perceive as a non-causal pass. The duration of one such event was 175 ms. All events moved from left to right. In addition to the event in the test location, which was located 1.5 dva below fixation, on two thirds of the trials, we presented an additional event at the context location (1.5 dva above fixation). This context event, when present, was either a launch (0% overlap), or a pass (100% overlap). Participants were instructed to report their perception (i.e., launch vs. pass) for the event presented below fixation. During this process, observers had to maintain fixation at the center of the screen. Trials would start once observers successfully maintained fixation for at least 200 ms. Each block consisted of 42 trials (7 disk overlaps × 3 context types × 2 repetitions). The first session was used as a training session and featured only 10 blocks (all without adaptation), while the two test sessions featured 16 blocks each (8 without adaptation; 8 with adaptation).

The first 8 blocks of the two test sessions were without adaptation to determine the perception of causality before adaptation (**Figure 1b**). After the 8th block, each block started with an adaptation sequence of 320 launch events presented in quick succession. The direction of each single event in the launch adaptor was randomly chosen from a narrow uniform distribution around the direction on the horizontal meridian (±30 degrees). After the adaptation sequence, each trial included a top-up adaptation of 16 events and then continued as before with no further changes. The only difference between the second and third session was the adaptation location, which could either be at the test event location or at the context event location, the order of which was randomly assigned across participants.

In all experiments, we tracked eye movements to ensure proper fixation behavior during presentation of the test events and presentation of the adaptors. More specifically, we tracked the dominant eye’s current position at a sampling rate of 1000 Hz and determined online the eyes’ distance to the screen center. We aborted a trial, whenever the distance between eye position and screen center exceeded 2 dva. Observers repeated these trials at the end of a block in randomized order. During presentation of the adaptors, trials were not aborted when observers broke fixation. Instead, we presented a short message (at the fixation point) asking observers to please fixate in the center of the screen once the observers’ gaze exceeded 2 dva away from the screen center.

### Data analysis

We estimated psychometric functions relating disk overlap to the proportion of reported launches using logistic functions with four parameters for the intercept, slope, as well as upper and lower asymptotes. We fitted these functions separately for each observer and condition. However, we did not obtain points of subjective equality (PSE) for each observer and each condition, as some participants’ proportion of reported launches was below 0.5 in some conditions. For inferential statistics, we therefore determined the proportion of reported launches for each disk overlap and condition, computed the difference between two conditions of interest, and obtained the cumulative sum across the disk overlaps (see Results for details). A significant interaction in the rmANOVA was complemented by running post-hoc paired t-tests. In all figures, error bars indicate ±1 within-subject standard error of the mean (SEM; Baguley, 2012; Morey, 2008).

The data and all original code have been deposited at the Open Science Framework and is publicly available as of the date of publication. [LINK].

## Results

We quantified the perception of causality by determining the proportion of reported launches for test events with varying disk overlap, ranging from clear launches (0% overlap) to clear passes (100% overlap). We determined whether additional context events would bias the report for ambiguous test events in the direction of the context event. We then asked whether interfering with the local launch detection mechanism at the location of the test event by means of visual adaptation would attenuate, or even eliminate, contextual influences.

### Strong aftereffects from adaptation only in trials without context

Visual adaptation successfully affected the perception of causality in no-context trials (**Figure 2a–b**), replicating previous findings (Kominsky & Scholl, 2020; Ohl & Rolfs, 2025; Rolfs et al., 2013): Observers showed a strong negative aftereffect in that they were less likely to report a causal launch after adaptation as compared to before adaptation. Notably, in context-trials, we did not observe a change in the proportion of reported launches from before to after adaptation. This result was corroborated by a one-way (3 context types: no-context vs. launch-context vs. pass-context) repeated-measures analysis of variance (rmANOVA) in which we assessed the magnitude of adaptation in each condition. To this end, we first subtracted the proportion of reported launches before adaptation from that after adaptation (**Figure 2b**). We then determined the cumulative adaptation (ca) across all disk overlaps. When ca is significantly smaller than 0, it would confirm a reliable reduction in perceived causality, that is, a negative aftereffect. The analysis revealed a significant main effect of context type (F(2, 18) = 10.09, p = 0.001). Post-hoc t-tests confirmed a significant adaptation in no-context trials (t(9) = –6.06, p < 0.001, ca_no_context_ = –1.15, CI_95%_ = [–1.58, –0.72]), but not in trials with launch context (t(9) = 0.08, p > 0.250, ca_launch_context_ = 0.02, CI_95%_ = [–0.54, 0.58]), or pass context (t(9) = –1.18, p > 0.250, ca_pass_context_ = –0.29, CI_95%_ = [–0.85, 0.27]). Moreover, adaptation was stronger in no-context trials as compared to launch-context trials (t(9) = –5.39, p < 0.001, Δca = –1.17, CI_95%_ = [–1.66, –0.68]) and pass-context trials (t(9) = –2.63, p = 0.027, Δca = –0.86, CI_95%_ = [–1.59, –0.12]). The impact of adaptation was not statistically different between the launch context and pass context (t(9) = 1.23, p > 0.250, Δca = 0.31, CI_95%_ = [–0.26, 0.89]).

**Figure 2.**
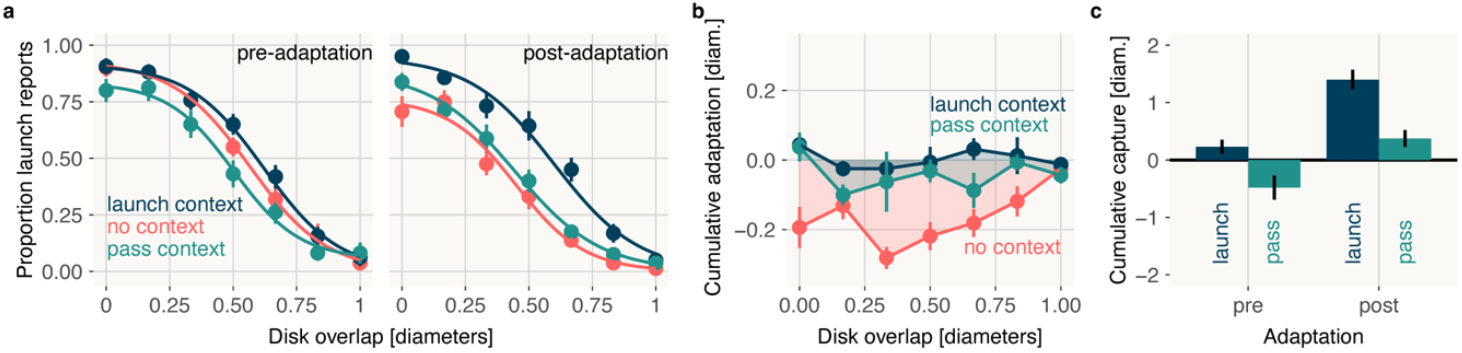
Adaptation at the test location. **a** Mean proportion of launch reports as a function of disk overlap, plotted separately for each context event (no context in red, launch context in blue, pass context in green) and displayed separately in blocks before adaptation (left panel) and after adaptation (right panel). Visualization of psychometric curves is based on fitting the model parameters to the mean reported launches in an experimental condition. **b** The magnitude of adaptation obtained for the three context conditions calculated by subtracting the mean proportion of reported launches before adaptation from the mean proportion of reported launches after adaptation. **c** Cumulative context effects (computed separately for pre-vs. post-adaptation and for launch vs. pass contexts) are determined as the differences in proportions of reported launches in the no context condition and the context conditions, accumulated over all disk overlaps. Error bars are ± 1 SEM.

### Strong context effects after visual adaptation

Context events changed observers’ proportion of causal reports in the predicted direction. Before adaptation, observers were less likely to report a causal launch when a simultaneous pass context event was presented. Similarly, they reported more causal launches in the presence of a launch context event (i.e., causal capture; Choi & Scholl, 2004, 2006; Scholl & Nakayama, 2002; **Figure 2a**, left panel). After adaptation, we found a strong launch context effect in the different psychometric curves for no-context trials and launch-context trials (**Figure 2a**, right panel).

We quantified the magnitude of the contextual influence by subtracting the proportion of launch reports in no-context trials from context trials for each disk overlap. In a second step, we then computed the cumulative context effect over the disk overlaps. Positive values, thus, indicate more causal launch reports in the presence of a context event, while negative values show that observers reported fewer launches (i.e., more passes).

Before adaptation, we observed a small positive launch context effect and a small negative pass context effect (**Figure 2c**). However, after adaptation, the context effect was strongly increased for the launch context. We corroborated this observation by a two-way (adaptation: before vs. after; context type: launch-context vs. pass-context) rmANOVA. The analysis revealed a significant main effect of context type (F(1, 9) = 10.47, p = 0.010): Observers reported more launch reports when the launch-context as compared to the pass-context was presented. After adaptation, we observed a significant increase in the proportion of reported launches (F(1, 9) = 17.07, p = 0.003), that is, more launch reports in both context conditions as compared to no-context trials. That finding can be reconciled with the results above that showed a strong negative aftereffect following adaptation in no-context trials and yet no influence of adaptation in context trials. Consequently, the influence of the launch context increased strongly after visual adaptation, effectively rescinding the negative aftereffect.

### Visual adaptation is spatially specific

In an additional session, we presented an adaptor at the context event location. In the no-context condition, this allowed us to determine the spatial specificity of adaptation in our experimental setup where test and context events were separated by 3 dva. More specifically, we assessed whether an adaptor presented at one location would also elicit adaptation at the other relevant location. Moreover, we predicted that adaptation at the context location would reduce a launch context’s (but not a pass context’s) impact on the perception of causality at the test location

This condition ruled out a spatially broadly tuned adaptation effect. The adaptor at the context event location did not result in adaptation for test events in no-context trials (**Figure 3a-b**). This observation was corroborated by a one-way (3 context types: no-context vs. launch-context vs. pass-context) rmANOVA in which we again quantified the magnitude of adaptation using the cumulative adaptation score introduced above. We did not observe statistically significant adaptation when the adaptor was presented at the context location (F(2, 18) = 2.82, p = 0.09). Critically, the adaptation was not significant in any context condition, including no-context trials, (t(9) = –1.57, p = 0.151, ca = –0.24, CI_95%_ = [–0.58, 0.10]), launch-context trials (t(9) = –0.41, p > 0.250, ca = –0.08, CI_95%_ = [–0.53, 0.36]), and pass-context trials (t(9) = 1.37, p = 0.204, ca = 0.23, CI_95%_ = [–0.15, 0.60]).

**Figure 3.**
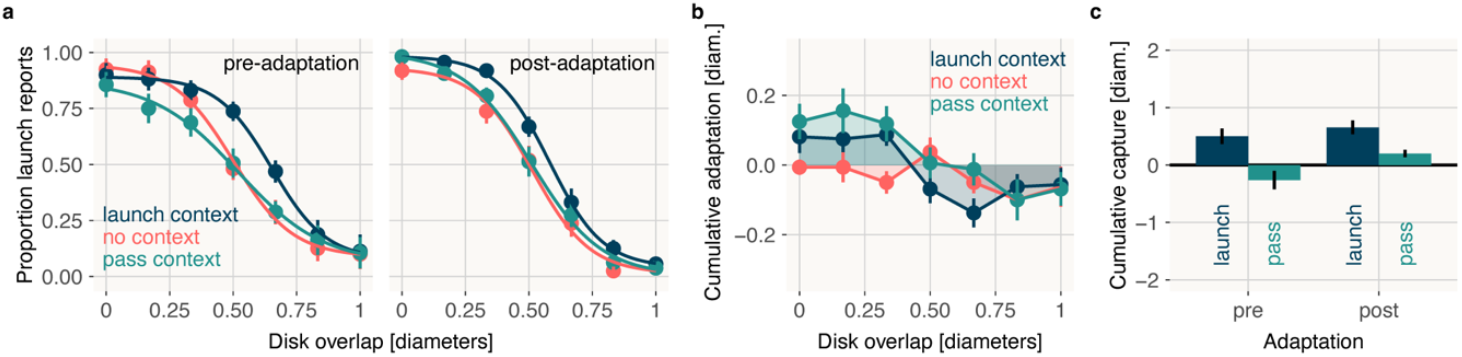
Adaptation at the context location. **a** Mean proportion of launch reports as a function of disk overlap and as a function of the context event (no context trials in red, launch context in blue, pass context in green) displayed separately in blocks before adaptation (left panel) and after adaptation (right panel). Visualization of psychometric curves is based on fitting the model parameters to the mean reported launches in an experimental condition. **b** The magnitude of adaptation obtained for the three context conditions (no context in red, launch context in blue, pass context in green) by subtracting the mean reported launches before adaptation from the mean reported launches after adaptation. **c** Cumulative context effects (computed separately for pre-vs. post-adaptation and for launch vs. pass contexts) are determined as the differences in proportions of reported launches in the no context condition and the context conditions, accumulated over all disk overlaps. Error bars are ± 1 SEM.

Moreover, we predicted that the influence of the context event may be attenuated after the adaptation at the context event location. Before adaptation, we observed a contextual influence of launches in the expected direction (**Figure 3c**). Observers reported more launches in ambiguous test events when the launch context event was presented. At the same time, we observed a trend that observers reported more passes in the presence of the pass-context event. Contrary to our expectations, adaptation at the context location did not alter this contextual influence. We still found that observers reported more launches when the launch-context was presented and there was no evidence for a contextual influence of the pass-context event. We corroborated these observations by a two-way (adaptation: before vs. after; context type: launch-context vs. pass-context) rmANOVA. The analysis revealed a significant main effect of context type (F(1, 9) = 7.96, p = 0.020): Observers reported to perceive more launches in the presence of a launch-context. Adaptation did not significantly change the contextual influence (F(1, 9) = 3.54, p = 0.092). Moreover, the interaction between adaptation and context type was not significant (F(1, 9) = 2.22, p = 0.171).

## Discussion

Using visual adaptation, we revealed that the contextual influence on the perception of causality does not depend on an unadapted launch detector at the test location. We successfully adapted the perception of causality at the test location, but the adaptation did not attenuate the contextual influence. This shows that the adapted launch detector is not sufficient to eliminate the influence of context events. We suggest that the contextual influence on the perception of causality bypasses the adapted launch detector and counteracts the visual adaptation.

Surprisingly, causal capture (i.e., the launch context effect) was strongest after adaptation at the test event location. Because we quantified the magnitude of causal capture by taking the difference in reported launches between no-context and launch-context trials, two results jointly accounted for this finding: While adaptation strongly influenced the reports in no-context trials, we did not observe any influence of adaptation in context trials.

Before adaptation, the context effects were considerably weaker than would be expected from previous studies (e.g., Scholl & Nakayama, 2002). How can we explain this discrepancy? There are several differences in the exact visual stimulus, all of which may have contributed to the weaker context effect (before adaptation) observed in our experiment. First, the two disks involved in the events were identical, whereas the stimuli in previous studies on causal capture were of different colors (Choi & Scholl, 2004, 2006; Scholl & Nakayama, 2002). A pass event consisting of two disks of different colors might more easily result in a launch percept given the disks’ distinct appearance. Moreover, we presented the test event below fixation and the context event above fixation at an equal vertical distance from the central fixation point. Observers presumably had strong evidence for perceiving a non-causal pass at the test event location. An additional context event may not have been able to overcome the strong perceptual evidence. In previous studies, the evidence at the test event location was reduced by either presenting the critical test event in the periphery or by presenting ambiguous test events with partial overlap (Choi & Scholl, 2004). While this evidence-based account provides an explanation for the small context effect before adaptation, it does not explain why causal perception in context trials after adaptation was the same as compared to before adaptation. Decreasing the perceptual evidence at the test event location, did neither result in more reported launches in launch-context trials nor fewer reported launches in pass-context trials. The reports in context trials were independent of the adaptation at the test event location. This finding suggests that the mechanism driving the context effects bypasses the adapted launch detector.

What mechanism can mediate the contextual influence and bypass the mechanism for launch detection? Contextual influences can be observed even in the absence of a launch-context event by simply presenting a spatially aligned group of stimuli that begins moving along with the second disk (Choi & Scholl, 2004). Increasing the grouping between the second disk of the test event and a context disk by means of Gestalt laws such as connectedness, good continuation, proximity, and common motion increased the contextual influence (Choi & Scholl, 2004). These results suggest that groups, in addition to single objects, constitute a unit over which a detection mechanism can operate to perceive launches. We suggest, therefore, as a possible explanation for the current findings that the visual adaptation only affected the perception of spatially specific causal interactions between individual objects. Context events, however, allowed the formation of a group consisting of the test and the context event. The detection of a causal relation at the level of the grouped stimuli then escaped the spatially specific adaptation. This explanation is also in line with our result that adaptation at the context event location did not decrease the contextual influence. Here again, grouping the context and test event together may have allowed the visual system to detect launches over parts of the group that were unaffected by the visual adaptation. Future studies should identify how exactly causal perception between individual objects and groups of objects relate to each other.

## Conclusion

We capitalized on the visual adaptation of causal perception as a powerful tool to uncover the rules underlying the detection of causal relations in our environment. Here we assessed the architecture underlying contextual influences on the perception of causal launches. We asked whether context events can exert their influence on other locations even if adaptation has reduced the propensity to detect launches at that location. Our findings revealed that visual adaptation strongly reduced causal perception at the test event location, but only in the absence of context events. In the presence of context events, context effects overruled the strong influence of visual adaptation, such that perceptual reports in context trials remained effectively unaffected by adaptation. In line with previous suggestions, we argue that context effects operate on a group of stimuli which allowed the contextual influences reported here to bypass the adapted test event location.

## Acknowledgments

This research was supported by a DFG research grant to S.O (OH 274/4-1) as well as funding from the Heisenberg Programme of the DFG to S.O. (OH 274/5-1). The authors declare no competing financial interests.

